# Retrograde tracing of breast cancer-associated sensory neurons

**DOI:** 10.1101/2024.02.26.582088

**Authors:** Svetllana Kallogjerovic, Inés Velázquez-Quesada, Rutva Hadap, Bojana Gligorijevic

## Abstract

Breast cancer is one of the leading causes of mortality among women. The tumor microenvironment, consisting of host cells and extracellular matrix, has been increasingly studied for its interplay with cancer cells, and the resulting effect on tumor progression. While the breast is one of the most innervated organs in the body, the role of neurons, and specifically sensory neurons, has been understudied, mostly for technical reasons. One of the reasons is the anatomy of sensory neurons: sensory neuron somas are located in the spine, and their axons can extend longer than a meter across the body to provide innervation in the breast. Next, neurons are challenging to culture, and there are no cell lines adequately representing the diversity of sensory neurons. Finally, sensory neurons are responsible for transporting several different types of signals to the brain, and there are many different subtypes of sensory neurons. The subtypes of sensory neurons which innervate and interact with breast tumors are unknown. To establish the tools for labeling and subtyping neurons that interact with breast cancer cells, we utilized two retrograde tracer’s standards in neuroscience, wheat-germ agglutinin (WGA) and cholera toxin subunit B (CTB). *In vitro*, we employed primary sensory neurons isolated from mouse dorsal root ganglia, cultured in a custom-built microfluidic device DACIT, that mimics the anatomical compartmentalization of the sensory neuron’s soma and axons. *In vivo*, we utilized both syngeneic and transgenic mouse models of mammary carcinoma. We show that CTB and WGA trace different but overlapping sensory neuronal subpopulations: while WGA is more efficient in labeling CGRP+ neurons, CTB is superior in labeling the NF200+ neurons. Surprisingly, both tracers are also taken up by a significant population of breast cancer cells, both *in vitro* and *in vivo*. In summary, we have established methodologies for retrograde tracing of sensory neurons interacting with breast cancer cells. Our tools will be useful for future studies of breast tumor innervation, and development of therapies targeting breast cancer-associated neuron subpopulations of sensory neurons.

## INTRODUCTION

The human nervous system is composed of the central and peripheral nervous system (CNS and PNS). With a well-defined structure and function, CNS and PNS interact to detect and respond to signals^1^. The PNS is composed of sympathetic, parasympathetic, motor, and sensory neurons^2^. The sensory neuronal bodies (somas) reside in the dorsal root ganglia (DRG), located close to the spine, and in the trigeminal ganglion. Sensory neurons are a pseudo-unipolar type of neurons and extend in two directions-reaching out to innervate the periphery of the body and internal organs, and through the spinal cord towards the brain^3^. Axons of sensory neurons are commonly grouped in bundles called fascicles and surrounded by extracellular matrix^4^. At the terminal end of axons are receptors specialized to detect various sensory signals. Sensory neurons can be classified based on the type of the stimulus that they sense, the axon thickness, the electrical impulse transmission speed, and their expressed receptors. However, such classes can be only broadly defined due to the neuronal complexity and heterogeneity^5,6^.

While the role of tumor microenvironment in tumor cell behavior has been widely recognized in the last decade, the role of neurons in tumor progression and metastasis has been understudied until recently. An increasing number of studies has reported that tumor innervation plays a significant role in the progression of different tumor types, including breast^7–10^. Specifically, the presence of thicker nerves in breast cancer patients has been associated with higher malignancy and worse prognosis^11^. Additionally, the presence of sensory neurons was shown to correlate with shorter survival in triple-negative breast cancer patients^12^. However, which subpopulations of neurons are mostly involved in breast tumor innervation and whether they are also present in healthy breast, is unknown.

Most *in vitro* studies studying host-cancer cell interactions are based on co-culturing techniques. However, this approach is not ideal in replicating the interaction between the sensory neurons and breast cancer cells. Due to the unique anatomy of the sensory neurons, microenvironments surrounding the soma and hence neurotransmitters secreted by the soma in the DRG and axons in the breast are very different^13,14^. To address this, we custom-designed a microfabricated device for axon-cancer cell interaction testing (DACIT)^15^, that compartmentalizes the soma and axonal terminals, separating them by microgrooves that guide axons between compartments.

Neuroscience has long employed the natural ability of some proteins to travel in retrograde direction towards deciphering neuronal connections in the CNS, and to a lesser extent, in PNS. A number of retrograde tracing approaches are available, including enzymatic tracers, such as horseradish peroxidase (HRP), toxin-based tracers, including tetanus or cholera toxin subunit B (CTB); plant-based lectins, including wheat germ agglutinin (WGA), and dyes such as Diamidino Yellow, Fast-blue, or Fluorogold (FG)^16–18^, to name a few^16^. Although dye-based tracers provide robust labeling, they come with a relatively wide emission spectrum, and can be toxic, non-selective, or impair neuronal integrity^17, 18^. As a result, most commonly used tracers are WGA and CTB. These tracers are taken up via the same general mechanism of receptor-ligand binding, however, CTB binds to the membrane glycan GM1 ganglioside, a sialic-acid containing glycosphingolipid^19^, whereas WGA binds to membrane receptor sugar N-acetylglucosamine, with a preference to dimers and trimers, and to N-acetylneuraminic (sialic) acid^20^. For example, in PNS, CTB has been used for tracing bladder^21^, liver^22^, mammary gland^23^ and pancreas innervation^22^, while WGA has been recently used to trace sensory neurons in the mouse model of ovarian cancer^24^.

To establish techniques for retrograde tracing of breast-cancer associated sensory neurons, we utilized WGA and CTB tracers in *in vitro* and *in vivo* conditions, tracing sensory neurons in the presence of breast cancer cells (BCCs). Our data demonstrated that while CTB was more applicable for tracing the NF200+ neurons, WGA was more efficient in tracing the CGRP+ neurons. Furthermore, our data revealed the tendency of both tracers to label cancer cells. Overall, we here present methodologies to retrograde trace and subtype sensory neurons that have axons in contact with breast cancer cells.

## MATERIALS AND METHODS

### Fabrication of the DACIT

DACIT devices were fabricated as previously described^15^. Briefly, a 10:1 ratio of Sylgard 184 elastomer base and curing agent (Electron Microscopy Sciences) were mixed and poured into a custom-fabricated SU-8 master. A desiccator (Nalgene, Thermo Fischer Scientific) was used to remove bubbles. The mixture was baked at 65°C overnight. The cured PDMS was carefully removed and cut out into 23 mm-squares. Glass coverslips (24 mm squares, #1405-10, Globe Scientific Inc) and the PDMS were bonded using a plasma machine (PE-25 Series Plasma System, Plasma Etch), and sterilized by ethanol and UV light. Before use, devices were coated overnight with 50 μg/ml Poly-L-lysine (Sigma-Aldrich) followed by 1:20 Matrigel in PBS (Corning).

### Mouse models

All mice experiments were conducted following NIH regulations and approved by Temple University IACUC. Experiments were performed on female mice, 7-10 weeks of age (Jackson Lab): Balb/cJ mice (#000651), C57BL/6J (#000664), B6.Cg-Tg(Thy1-YFP)16Jrs/J (#003709), FVB/NT (#001800), and FVB/N-Tg(MMTV-PyVT)634Mul/J (#002374). For syngeneic tumor models, 10^6^ of E0771-tdTomato cells resuspended in 100 μl PBS were injected in aseptic conditions into the 4^th^ mammary gland of mice.

### DRG recovery, dissociation, and plating

Primary Mouse Sensory Neurons (PMSN) were collected from healthy female, 7-8 weeks-old Balb/cJ mice immediately post-sacrifice. The spine was recovered and cleaned from excessive tissue. The vertebrae body and the spinal cord were removed, and the rest of the spine was cut in half in the sagittal plane to achieve better visualization of the DRGs found in the intervertebral neural foramen. The DRGs were immersed in 10 mM HEPES (Gibco) solution in HBSS Ca^+^, Mg^2+^-free (Gibco), on ice, and transferred to 5 mg/ml collagenase P (Sigma-Aldrich) for 45 minutes at 37°C, followed by 0.05 % Trypsin-EDTA (Gibco) for 4 minutes. The trypsin was inactivated by adding horse serum (Atlanta) in Neurobasal^TM^ medium (NB, Corning) supplemented with 2% B-27™ supplement, serum-free (Gibco) and penicillin-streptomycin (Gibco). Cell disaggregation was done with a polished pipet and the cell suspension was filtered with 40 μm cell strainer (Corning). Non-neuron cells were removed by centrifugating the cell suspension on Bovine Serum Albumin (Sigma-Aldrich) 10%. The concentrated neurons were then resuspended into a complete NB medium, counted using Trypan Blue and 7 x 10^3^ PMSNs were loaded into the left, neuronal compartment of DACIT, resuspended in 8 μl of NB complete media and incubated at 37°C for 25 minutes. Next, 200 μl of complete NB medium was added to the neuronal compartment, and 150 μl was added to the right, axonal compartment. One day (24 hours) after plating, the media was changed, and supplemented 1:2000 with Neuron Growth Factor (Ngf 2.5S Native Mouse Protein, Invitrogen) in the right, axonal compartment of the device.

### Cell culture

The tdTomato-expressing (Addgene, Cat# 54642) cell line E0771 (kind gift of Lucia Borriello in Condeelis Lab) was established using Lipofectamine 3000 transfection (Invitrogen™ L3000015), followed by 4-week of selection with 1 mg/ml G-418 (Research products international) and sorting of top 5% expressors. Stable culture of tdTomato-E0771 was maintained at 37 °C and 5% CO2, in DMEM (Corning), supplemented with 10% FBS (Atlanta Biologicals), 1 mg/ml G-418 and 1% penicillin – streptomycin (Gibco). Cells (2.5 x 10^3^ cells in 10 μl) were plated in axonal compartment of DACIT, followed by addition of 140 μl of DMEM 30 min later.

### Retrograde tracers

Retrograde tracers, CTB pre-labeled with Alexa 647, CTB-647 (Invitrogen) and WGA-488 (Invitrogen), were dissolved in PBS at 1mg/ml and loaded into DACIT or injected on mice. In DACIT, 100 μl of media was collected from the axonal compartment, mixed with 8 μl of tracer (1mg/ml) and then added back, assuring proper mixing. In mice, a total of 9 μl of tracer (1mg/ml) was injected in 3 sites surrounding the nipple (3μl/ injection), using a 10 µL, 26-gauge Hamilton syringe (#701 N SYR, Cemented NDL, 3 in, point style 2). The mammary fat pads, tumors, and DRGs were recovered 3-5 days later.

### Fixing, sectioning, and immunofluorescent labeling

Samples were fixed in 4% paraformaldehyde (Alpha Aesar) for 10 min at RT (PMSNs), 90 min at RT (DRGs), or 48 h at 4°C (mammary fat pad or tumor). The fixed DRGs were either labeled intact, embedded in 2% low-melting agarose (BP165-25, Fisher Bioreagents) prior to vibratome sectioning at 200 μm thickness (Vibratome Series 1000, Sectioning system), or frozen at –80 °C in optimal cutting temperature gel (OCT) for cryosectioning at 16 μm (Leica CM3050S). Permeabilization and blocking were done in PBS with 5% normal goat serum (NGS) and 1% Triton X-100 for 45 min at RT (PMSNs), or 48 h at 4°C (DRGs), followed by overnight incubation (PMSNs) or 96 h (DRGs) at 4°C with the mouse monoclonal anti-CGRP (1:200, Novus Biologicals, Cat# NB036540), chicken polyclonal to Neurofilament heavy polypeptide NF200 (1:250, Abcam, Cat# ab4680), rabbit polyclonal PGP9.5 (1:250, Abcam, Cat# ab15503), in 1% Triton X-100. Washing was performed with 0.05% Tween-20 in PBS, and neurons were incubated at RT for 45 min (PMSNs) or 48h (DRGs) in secondary goat antibodies: anti-mouse Alexa Fluor 405 (1:400, Invitrogen, A-31553), anti-chicken Alexa Fluor 488 (1:400, Abcam, ab150169), anti-chicken Alexa Fluor 647 (1:400, Invitrogen, A-21449), and with IB4-568 lectin (1:400, Invitrogen™ I21413).

### Imaging

Images were acquired by laser scanning hybrid confocal-multiphoton microscope (FV1200, Olympus), using a 10X dry (UPLXAPO10X, 0.4 NA, Olympus), 20X dry (UCPLFLN20X, 0.7 NA, Olympus), or 30X silicone oil immersion objective (UPLSAPO30XSIR, 1.05 NA, Olympus) with 1-5 µm z sections. Live imaging of the DACIT device was performed daily using an environmental chamber (Tokai Hit). The intact DRGs were covered with 20-40 μl of PBS and immobilized with a coverslip in a Mattek dish.

### Analysis

The images were processed using Fiji software. Five fields of view (FOV,. 18 x. 64 μm each) were followed in DACIT over 5 days to quantify the labeled somas and cancer cells. Cells were defined as positive when the signal over background ratio (S/N) was >3. Immunolabeled cells were reported when positive for antibody and co-localizing with CTB or WGA tracer. The single– and double-marked somas were expressed as a percentage of the CTB+ or WGA+ totals. The intact-DRGs labeled with PGP9.5+, were reported as the percentage of PGP9.5+ somas that are CTB+, WGA+, or CTB/WGA+.

## RESULTS

### Retrograde tracing of primary mouse sensory neurons using WGA or CTB

To test the ability of WGA and CTB to label primary mouse sensory neurons (PMSNs) *in vitro*, we employed a custom-build device for axon and cancer cell interaction testing (DACIT)^15^. Briefly, DACIT allows the two neuronal compartments, axons and somas, to be separated by microgrooves and cultured in different chambers of the device, under different conditions which mimic either DRG (left compartment, soma) or tumor environment (right compartment, axons). Freshly isolated neurons were plated in the soma compartment of DACIT at day *in vitro* 0 (DIV 0). Retrograde tracers CTB or WGA, were added to the axonal compartment on DIV 3, after confirming axons have successfully crossed the microgrooves and populated the axon compartment (**Figure 1A**). To track the increase in the number of labeled somas, we collected images daily, DIV 4-7 (**Figure 1B**). Our analysis revealed a sharp increase in the number of labeled soma between DIV 4 and DIV 5, followed by a slow and steady increase between DIV 5-7, a pattern observed in both tracers (**Figure 1C**). Interestingly, CTB consistently labeled a higher number of somas. We hypothesized that a reason may be that the subtypes of sensory neurons labeled by CTB are prevalent, and possibly different to those labeled by WGA. To test this, following daily images, cultures were fixed on DIV 8 and immunolabeled utilizing markers for three major subtypes of sensory neurons. This inclides NF200, a 200 kDa neurofilament marker that is present mainly in fast-responding myelinated Aβ and Aδ fibers, nociceptors responsible for pain, mechanoreceptors sensing touch and pressure, and proprioceptors, sensing movement and position^25^; calcitonin gene-related peptide (CGRP), which labels slow-responding, peptidergic thin C-fibers, and a sub-population of Aδ fibers^25,26^; and isolectin-B4 (IB4), that labels slow responders, small non-peptidergic C-fibers, and neurofilament-poor neurons^25,27^. In **Figure 1D**, PMSNs traced by WGA prior to immunolabeling are shown, while **Figure 1E** shows PMSNs traced by CTB. Our analysis showed that the WGA+ or CTB+ neurons come from fairly different subpopulations (**Figure 1F**). For example, almost 80% of WGA+ PMSNs are CGRP+. Specifically, we found that from the total of neuron soma labeled with WGA, 30.1% were CGRP+ only, 30.1% were double positive, for CGRP+ and NF200+, and 26.88% were CGRP+ and IB4+. In addition, 8.6% were only IB4+, while 3.2% were only NF200+ and 1.1% were positive for both NF200+ and IB4+. In contrast, almost CTB+ soma were 70% NF200+, out of which 30% were only NF200+, while 39.62% were NF200+ and CGRP+. Meanwhile, 11.32% were CGRP+, 9.43% were IB4+ and were 9.43% CGRP+ and IB4+ (**Figure 1F**). In summary, this indicates that WGA more efficiently traces CGRP+ neurons, while CTB is more efficient in NF200+ neurons.

**Figure 1.**
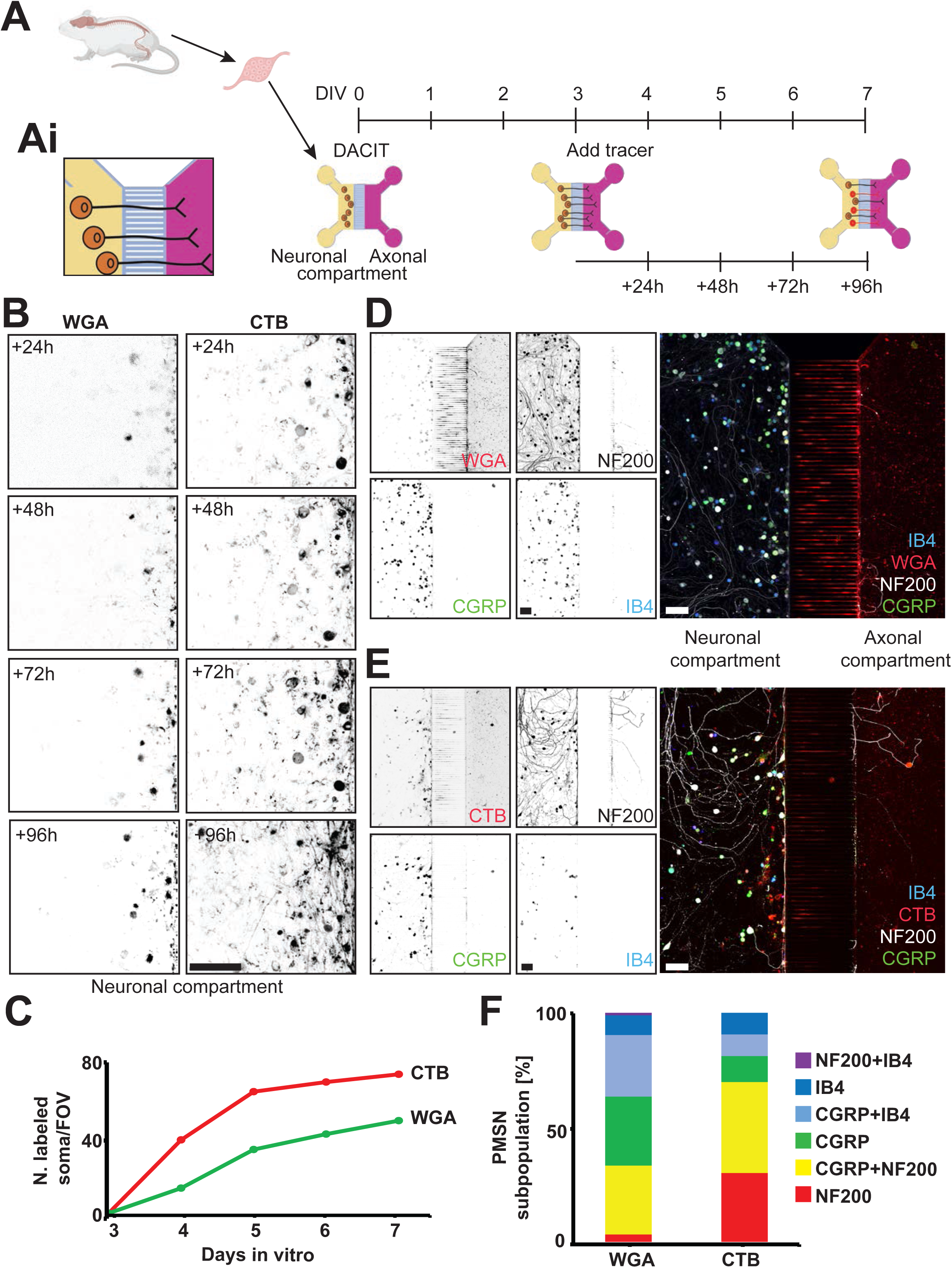
Retrograde tracing of sensory neurons *in vitro*. **A**. Steps for isolation and retrograde tracing of PMSNs: Mouse DRGs are dissociated and retrieved PMSNs are loaded into the neuronal compartment (yellow, left) of DACIT at DIV 0. At DIV 3, when axons cross the microgrooves and reach the axonal compartment (pink, right), the tracers are added to the axonal compartment. Images are collected daily DIV 4-7 and PMSNs are fixed and immunolabeled on DIV 8. Insert **Ai**. A zoomed-in image of the DACIT showing the FOV depicted in D and E. Figure created in Biorender. **B.** Representative images of the neuronal compartment of DACIT at 24-96h (DIV 4-7), showing PMSN somas labeled via retrograde tracing. Scale bar = 100µm. **C.** Number (N) of somas labeled by WGA (green line) or CTB (red line) over time. **D-E.** Representative images of immunofluorescence in the retrograde traced PMSN, either using WGA (**D**) or CTB (**E**). NF200 (grey) and CGRP (green) antibodies, and IB4 (blue). Scale bar: 100μm. **F.** Quantification of the WGA– or CTB-labeled PMSN subpopulations from **D-E**: NF200 (red) and CGRP (green) antibodies, and IB4 (blue). Colocalization of the markers is shown: CGRP and NF200 (yellow), CGRP and IB4 (cyan), and NF200 and IB4 (magenta).

### Retrograde tracing of sensory neurons in the presence of breast cancer cells

To test the ability of WGA and CTB to trace the PMSNs in the presence of BCCs, we loaded BCCs in the axonal compartment of the DACIT on DIV 2, and added tracers 24h later, on DIV 3 (**Figure 2A**), followed by daily imaging (**Figure 2B**). Here, similarly to **Figure 1C**, both tracers had a sharp increase in the total number of labeled soma DIV 3-5, followed by a slower increase DIV 5-7. We also analyzed co-labeled neurons, WGA+ and CTB+, confirming that only a small subpopulation of PMSN can be traced by either marker. (**Figure 2C**). Importantly, we also observed cancer cells labeling by tracers. While the rate of the WGA uptake was somewhat higher than CTB, both tracers labeled a similar number of cells at DIV 7 (**Figure 2D**).

**Figure 2.**
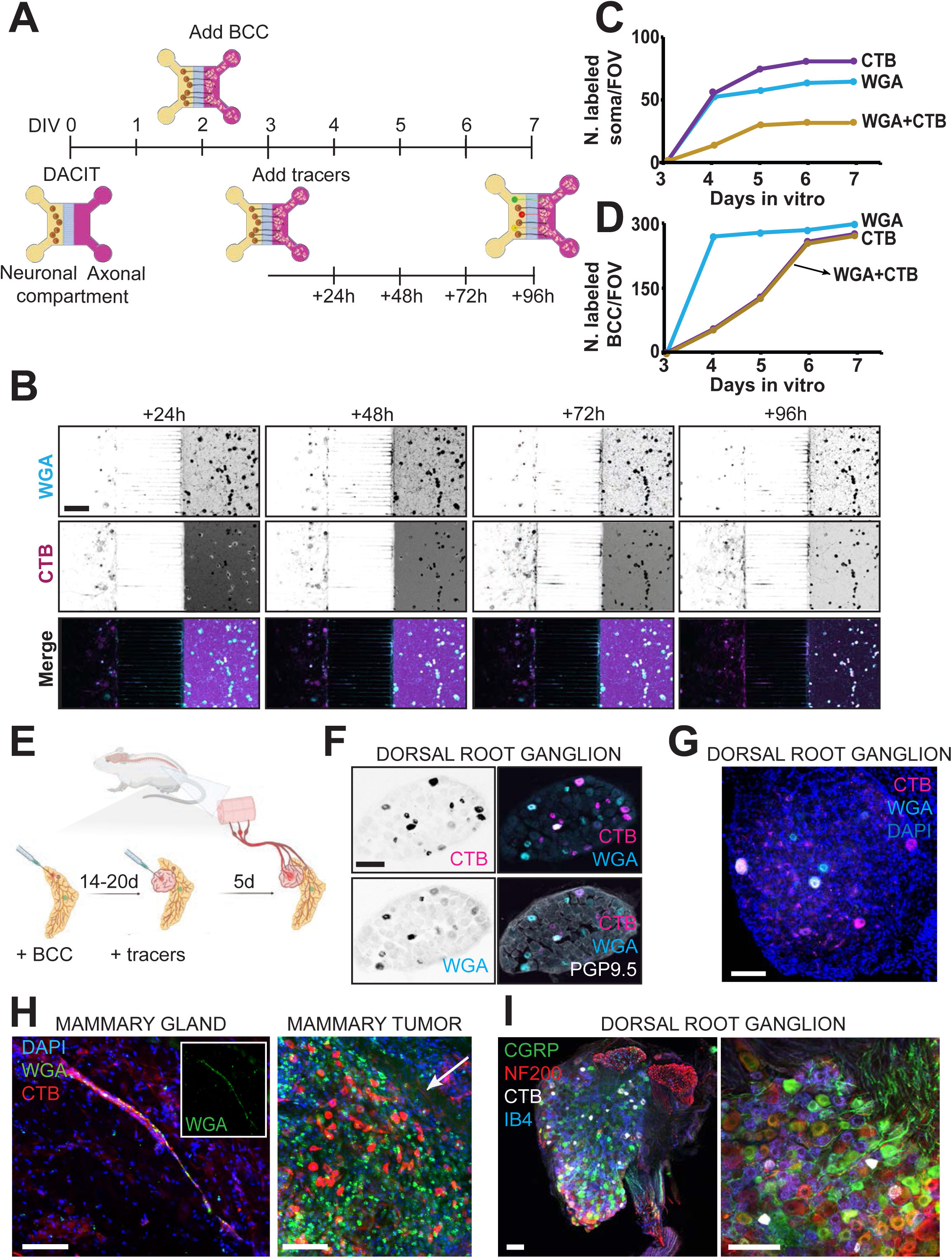
Retrograde tracing of sensory neurons in the presence of breast cancer cells. **A**. Steps for retrograde tracing of PMSNs in the presence of cancer cells: Sensory neurons are loaded into the neuronal compartment (yellow, left) of DACIT. At DIV 2, breast cancer cells are added to the axonal compartment (pink, right). At DIV 3, tracers are added to the axonal compartment. Images are collected daily DIV 4-7. Figure created in Biorender. **B.** Representative area of DACIT, showing somas in the neuronal compartment (left), axons extending along microgrooves (middle) and BCCs in the axonal compartment (right) at 24 – 96h (DIV 4-7). Scale bar = 100µm. **C.** Number of somas labeled by WGA (cyan), CTB (magenta), or both tracers (yellow line) over time. **D.** Number of BCCs labeled by WGA (cyan), CTB (magenta), or both tracers (yellow line) over time. **E.** Steps for retrograde tracing of the sensory neurons *in vivo*. Murine mammary cancer cells E0771-tdTomato are injected into the mammary fat pad of mice. When the tumor mass is palpable, tracers are injected into the healthy mammary fat pad or the mammary tumor. Five days later, DRGs are collected for analysis. Figure created in Biorender **F.** A cryosection of a DRG from a healthy mouse injected with WGA (cyan) and CTB (magenta) and stained with PGP9.5 (grey). Scale bar: 100µm. **G.** An intact DRG from a tumor-bearing C57BL/6J mouse injected with WGA (cyan) and CTB (magenta) and stained with DAPI (blue). Scale bar: 100µm. **H.** Mammary fat pad (left) and tumor (right) injected with WGA (green) and CTB (red) and stained with DAPI (blue). Arrow pointing at CTB+ axon (red). Scale bar: 100µm. **I.** A DRG vibratome section from a tumor-bearing mouse injected with CTB (red) and labeled with NF200 (grey), CGRP (green), IB4 (blue). Scale bar: 100 µm.

To test *in vivo* tracing, we utilized a syngeneic mammary tumor model. Cell line E0771-tdTomato was injected in the inguinal mammary fat pad of the C57BL/6J mice. When the tumor was palpable (20 days), WGA and CTB were injected into the healthy mammary fat pad or into the tumor (**Figure 2E**). Following a 5-day incubation, DRGs and tumors were collected (**Figure 2F-G**) and labeled somas were quantified. In healthy mice (**Figure 2F**), approximately 9% of the total PGP9.5+ somas were CTB+, or WGA+, and 1.17% of somas were co-labeled. In contrast, in tumor-bearing mice (**Figure 2G**), <1% were CTB+, or WGA+, suggesting a possible restricted diffusion, or uptake of tracer by non-neuronal cell types in tumors. To test if tracers were uptaken by host or cancer cells, we examined the localization of tracers in the mammary fat pad. We observed that CTB+ nerve fascicles (**Figure 2H**) contain only a few WGA+ axons (**Figure 2H** insert), confirming that in tumors, similar to *in vitro* conditions, only a small portion of tracers colocalize. Next, a portion of tracers was labeling BCCs, extracellular matrix and host cells. Finally, fixed and immunofluorescently labeled DRGs showed that CTB+ somas were either NF200+, CGRP+ or IB4+ (**Figure 2I**). Importantly, this result suggests that innervation of breast tumors involves NF200+, CGRP+ and IB4+ subpopulations of sensory neurons.

### Retrograde tracing of breast-cancer associated sensory neurons in the transgenic mouse models

To confirm and expand our results, we employed two transgenic mouse models: PyMT-MMTV, a model of mammary tumors in the FVB background (**Figure 3A**), and Thy1-EYFP-16 in C57/BL6J background, where neurons express fluorescent EYFP protein (**Figure 3B-E**).

**Figure 3.**
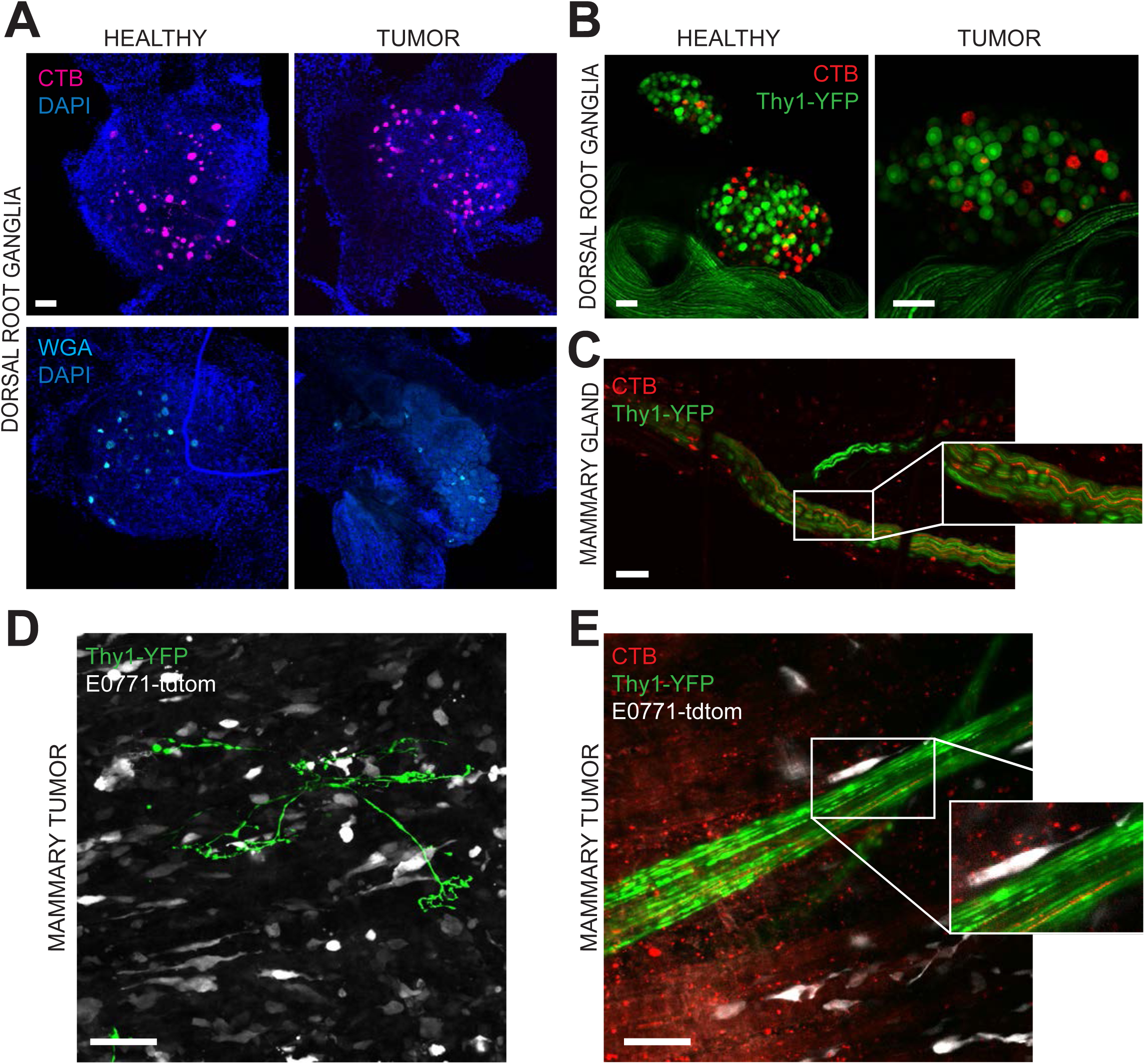
Retrograde tracing of breast-cancer associated sensory neurons in transgenic mice models. **A**. Intact DRGs from FVB mice (healthy, left; tumor-bearing PyMT-MMTV, right) injected with CTB (magenta, upper panel) or WGA (cyan, lower panel) and stained with DAPI (blue). Scale bar: 100μm **B.** An intact DRG from B6.Cg-Tg(Thy1-YFP)16Jrs/J transgenic mouse (YFP, green), injected with CTB (red) in the mammary fat pad (left) or in the E0771-tdTomato (grey) mammary tumor (right). Scale bars: 100μm. **C.** Intact mammary fat pad of the B6.Cg-Tg(Thy1-YFP)16Jrs/J mouse injected with CTB (red). Zoom-in shows CTB-labeled axons (red) within a nerve fascicle (YFP, green). **D.** Innervation of the E0771-tdTomato syngeneic mammary tumor (grey) in the B6.Cg-Tg(Thy1-YFP)16Jrs/J (YFP, green) mouse. Scale bar: 100µm. **E.** Mammary tumor (E0771-tdTomato, grey) traced via CTB (red) in the B6.Cg-Tg(Thy1-YFP)16Jrs/J (YFP, green) mouse. Zoom-in shows a single CTB-labeled axon within a nerve fascicle. Scale bar: 100µm.

First, retrograde tracing was tested in the PyMT-MMTV model which spontaneously forms mammary tumors. This model is characterized by an early onset of carcinoma, with a pattern of tumor progression similar to human breast cancer^28^. As all ten mouse mammary glands develop tumors during puberty, this allows simultaneous injection and further comparison of tracers in two sides of the inguinal mammary fat pad (in control FVB mice, left panels, **Figure 3A**), or mammary tumors (in PyMT-MMTV mice, right panels, **Figure 3A**). At 5 days post-injection, the DRGs were collected, confirming that injections in mammary fat pad, or mammary tumors, can be used to retrogradely trace sensory neurons innervating the breast in the transgenic mouse models.

To test CTB tracing in the animals with endogenously labeled neurons, we chose the B6.Cg-Tg(Thy1-YFP)16Jrs/J substrain of Thy1-YFP mice, which was reported to have the highest YFP expression in the sensory and motor neurons^29^. Sensory neuron tracing by CTB in the healthy mouse showed that most somas located in the DRG were YFP+, and a few of them were also CTB+ (**Figure 3B**, left). In the mammary gland, YFP-expressing fascicles also contained CTB+ axons (**Figure 3C**). To form syngeneic mammary tumors, we injected E0771-tdTomato cells in the inguinal mammary fat pad. Following retrograde tracing by CTB, we demonstrated that similarly to the healthy mouse, most DRG somas were YFP+, and some of them were also CTB+ (**Figure 3B**, right). Also, mammary tumor was densely innervated by bright Thy1-YFP+ axons (**Figure 3D**) and nerve fascicles which contain CTB+ axons (**Figure 3E**).

## DISCUSSION

In this study, we established methodologies to retrograde trace and subtype sensory neurons which have axons in contact with BCCs. Towards this goal, we utilized WGA and CTB retrograde tracers and immunofluorescence-based subtyping of sensory neurons in *in vitro* and *in vivo* settings. Our results suggest that while CTB is more efficient in tracing NF200+ fibers, WGA is more efficient in tracing CGRP+ C-fibers. Our data are consistent with a previous study which reported that GM1 was more abundant in NF200+ somas, compared to CGRP+ ones^30^. Additionally, our data showed that the co-localization of the two tracers occurred only in a small number of somas compared to each of the tracers individually, suggesting that the tracers should be used in a complementary fashion to trace tissues innervated by both NF200+ and CGRP+ subtypes of sensory neurons.

Our data demonstrates that both tracers strongly label breast cancer cells^31,32^, both *in vitro* and *in vivo*. Additionally, a small subpopulation of non-neuronal host cells in tumors were positive for tracers. This is possibly a result of the CTB’s ability to bind to immune cells, including B cells, dendritic cells and macrophages^33^, and WGA’s binding to the extracellular matrix and the membrane of the fibroblast^34^. While non-specific labeling of cancer cells does not negatively affect visualization of the retrograde-traced somas in DRG, it is important to keep in mind that a portion of the tracer cannot freely diffuse as it’s trapped by the tumor cells. Combined with the universal challenge of restricted diffusion in tumors due to the presence of cell-dense and matrix-dense regions, larger doses of tracers are required to achieve similar results in tumor-bearing, compared to the healthy animals.

To circumvent lack of specificity posed by conventional tracers, engineered viruses can be employed to target neurons, or specific neuron subpopulations. While many virus families have a natural ability to be internalized at the axonal terminals and travel along neurons in a retrograde fashion, including rabies viruses, herpes simplex and lentiviruses, many of them demonstrated limitations such as induced toxicity (rabies) or low level of expression (lentiviruses)^35^. To address these limitations, retrograde-Adeno-Associated Viruses (AAV-retro) were recently developed^35,36^. By combining fluorescent protein-encoding AAV-retros with serotypes with highest tropism for sensory neurons (AAV2, AAV9^37^), and synapsin promoter, most neurons innervating tumors will be labeled, while CGRP or NF200 promoters would label specifically CGRP+ and NF200+ subpopulations^38,39^. In addition to retrograde tracing, AAV-retros can be used to silence or eliminate the targeted neurons.

Imaging of intact tissues allows the 3D reconstruction of the tumor innervation *in situ* and visualization of almost any tumor microenvironment components. However, probe penetration often poses challenges, more pronounced in immunofluorescence due to the size of antibodies, but present even with smaller labels, such as DAPI or fluorescent IB4-lectin used in our study. To address this, few groups have employed increased barometric pressure^40^ or electric field^41^ to improve antibody penetration. We have used both *in situ* labeled tissues, as well as vibratome– or cryotome-sectioning, followed by immunolabeling. In addition, we utilized Thy1-YFP mice with endogenously labeled neurons. Specific substrain used, B6.Cg-Tg(Thy1-YFP)16Jrs/J, was reported to label most sensory neurons, in addition to motor neurons and some of the neurons in the central nervous system^29^. Others have reported that while large myelinated sensory neurons and their axons are labeled brightly in this substrain, some of the TRPV1+ neurons are not labeled^42^. As our data shows almost no colocalization between CTB and Thy1-YFP, our preliminary conclusion is that the breast tumors may be highly innervated by CGRP+, small myelinated NF200+ neurons and potentially, TRPV1+ neurons, large myelinated sensory neurons are either absent, or present at a low density. In the future, a more comprehensive tracing of the immunolabelled intact tissue may be accomplished using tissue clearing followed by light sheet microscopy, which allows for 3D reconstruction of the entire mammary tumor or DRG. A recent study elegantly demonstrated that immunolabeling and clearing can be done using entire mouse via wild DISCO protocol^43^, which was also shown suitable for peripheral nervous system. With this approach, tumor innervation could be traced across the entire body, from breast tumor to DRG, and combined with sensory neuron immunolabeling.

In summary, our methods will be useful to study subpopulations of breast-cancer associated sensory neurons, likely affecting tumor growth and metastasis.

## ACKNOWLEDGMENTS

We are grateful to Professors Edna “Eti” Cukierman from Fox Chase Cancer Center, George Smith from Shriners Hospital at Lewis Katz Medical School of Temple University, Brian M. Davis at University of Pittsburgh for insightful discussions, and MS Elizaveta Belova from Gligorijevic Lab for her support during the study. Funding was provided by NIH NCI R01 CA230777; American Cancer Society Research Scholar Grant 134415-RSG-20-34-01-CSM, DOD BCRP BC230197 and WW Smith Charitable Trust, all to B.G.

## AUTHOR CONTRIBUTIONS

Conceptualization: S.K., I.V.Q, B.G., Data acquisition and analysis: S.K., R.H., B.G. Supervision: B.G., Writing: S.K., I.V.Q, B.G.

